# Network Topology Dynamics of Circulating Biomarkers and Cognitive Performance in Older Cytomegalovirus-seropositive or -seronegative Men and Women

**DOI:** 10.1101/673905

**Authors:** Svetlana Di Benedetto, Ludmila Müller, Stefanie Rauskolb, Michael Sendtner, Martin Fassnacht, Graham Pawelec, Viktor Müller

**Affiliations:** Max Planck Institute for Human Development, Berlin, Germany; University of Tübingen, Tübingen, Germany; Institute of Clinical Neurobiology, Würzburg, Germany; University Hospital, University of Würzburg, Würzburg, Germany

**Keywords:** aging, immunosenescence, inflammatory markers, cytokines, cytomegalovirus, neurotrophic and metabolic factors, cognition, network topology

## Abstract

**Background:** Cytokines are signaling molecules operating within complex cascade patterns and having exceptional modulatory functions. They impact various physiological processes such as neuroendocrine and metabolic interactions, neurotrophins’ metabolism, neuroplasticity, and may affect behavior and cognition. In our previous study, we found that sex and Cytomegalovirus (CMV)-serostatus may modulate levels of circulating pro- and anti-inflammatory cytokines, metabolic factors, immune cells, and cognitive performance, as well as associations between them.

**Results:** In the present study, we used a graph-theoretical approach to investigate the network topology dynamics of 22 circulating biomarkers and 11 measures of cognitive performance in 161 older participants recruited to undergo a six-months training intervention. For network construction, we applied coefficient of determination (*R*^2^) that was calculated for all possible pairs of variables (N = 33) in four groups (CMV^−^ men and women; CMV^+^ men and women). Network topology has been evaluated by clustering coefficient (*CC*) and characteristic path length (*CPL*) as well as local (*E*_*local*_) and global (*E*_*global*_) efficiency, showing the degree of network segregation (*CC* and *E*_*local*_) and integration (*CPL* and *E*_*global*_). We found that networks under consideration showed small-world networks properties with more random characteristics. Mean *CC*, as well as local and global efficiency were highest and *CPL* shortest in CMV^−^ males (having lowest inflammatory status and highest cognitive performance). CMV^−^ and CMV^+^ females did not show any significant differences. Modularity analyses showed that the networks exhibit in all cases highly differentiated modular organization (with *Q*-value ranged between 0.397 and 0.453).

**Conclusions:** In this work, we found that segregation and integration properties of the network were notably stronger in the group with balanced inflammatory status. We were also able to confirm our previous findings that CMV-infection and sex modulate multiple circulating biomarkers and cognitive performance and that balanced inflammatory and metabolic status in elderly contributes to better cognitive functioning. Thus, network analyses provide a useful strategy for visualization and quantitative description of multiple interactions between various circulating pro- and anti-inflammatory biomarkers, hormones, neurotrophic and metabolic factors, immune cells, and measures of cognitive performance and can be in general applied for analyzing interactions between different physiological systems.

## Introduction

Aging is accompanied by chronic low-grade inflammation that has been repeatedly identified even in overtly healthy individuals and is characterized by elevated levels of circulating pro-inflammatory cytokines (1). Cytokines represent signaling molecules having exceptional modulatory functions. They impact virtually every physiological process such as neurotransmitter metabolism, neuroendocrine interactions, and neuroplasticity, thereby not only affecting general health but also immunity and cognitive functioning (2–4). The cytokine network, containing cytokines, their receptors, and their regulators, is present in the brain and in various other physiological systems, and is highly controlled throughout the lifespan (5, 6). Cytokines and their receptors operate within multifactorial networks and may act synergistically or antagonistically in a time- and concentration-dependent patterns. These interactions allow cross-communication between different cell types, at different hierarchical levels, translating environmental signals into molecular signals (2, 7). The pro-inflammatory profile becomes strategic throughout the lifespan (8–11) - an increase of cytokine secretion, also thought to be associated with the influence of CMV-infection, may be at least partly responsible for age-associated degenerative disorders (12–16). Previous studies usually investigated individual roles of different cytokines, inflammatory mediators or metabolic factors in the age-related physiological alterations (17–21). With growing numbers of biomarkers, however, it may become difficult to interpret results and translate them into useful information.

In our recent work (22), we assessed inflammatory status and cognitive performance in 161 older participants recruited to undergo a six-month training intervention. We demonstrated that sex and CMV-latency have influence on levels of circulating pro- and anti-inflammatory cytokines, receptor antagonist, soluble receptor, metabolic factors, and immune cells. We also found that CMV-latency has modulatory effects on associations between individual peripheral biomarkers (22). Furthermore, we revealed an interaction between CMV-serostatus and sex associations with cognitive abilities: sex differences in fluid intelligence and working memory were noted only in CMV-negative individuals. Even more strikingly, the same group of elderly men also exhibited a lower inflammatory status in their peripheral circulation. Therefore, a well-balanced inflammatory and anti-inflammatory equilibrium appeared apparently to be decisive for optimal physiological functions and for optimal cognitive functioning.

Pro-inflammatory cytokines often act as negative regulatory signals modulating the action of hormones and neurotrophic factors. An unbalanced cytokine state may also affect the neuroendocrine system (and vice versa) impairing interplay between them, and contributing to disrupted homeostasis (23). Therefore, in the present study, we additionally considered such hormones as cortisol and dehydroepiandrosterone (DHEA) as well as neurotrophines and their regulators (insulin-like growth factor-1, IGF-1, and IGF-binding protein, IGFBP-3), to gain a more comprehensive image of these processes. Furthermore, we extended the number of inflammation-related metabolic factors and included measures of C-reactive protein (CRP) in our present analyses. Finally, instead of focusing on four latent factors representing the main cognitive abilities (as we did in the previous study), we included in our present analysis all 11 individual cognitive performance scores assessed within the cognitive battery of elderly individuals. Increasing complexity arose when attempting to analyze dynamic interconnections between all these factors and to investigate the modulatory impact of CMV-latency and sexual dimorphism. In an effort to better understand the relationships between the multiple circulating and functional biomarkers and to compare them regardless of their physiological hierarchical assignments, we applied a graph-theoretical approach and described constructed networks in terms of network topology and modular organization of network elements.

As stated by Bhavnani et al., network analyses offer two main advantages for studying complex physiological interactions: (i) they do not require a priori assumptions about the relationship of nodes within the data, such as the categorized assumption of hierarchical clustering; and (ii) they allow the simultaneous visualization of several raw values (such as cytokine or/and cell values, functional attributes), as well as aggregated values, and clusters in a uniform visual representation (24). This allows not only the more rapid generation of hypotheses based on complicated multivariate interactions, but also the validation, visualization, and confirmation of the results, obtained with other methodological approaches. Moreover, this enables a more informed methodology for selecting quantitative methods to compare the patterns obtained in the different sets of data regardless of their physiological hierarchical levels (24).

The purpose of the present study was to visualize and to quantitatively describe by means of a graph-theoretical approach the complex multiple interactions among diverse pro- and anti-inflammatory mediators, immune cell populations, hormones, neurotrophic and metabolic factors as well as cognitive performance in older CMV-seropositive and -negative men and women. Moreover, we aimed to design a new strategy for quantitative investigations of the network topology dynamics in circulating biomarkers and measures of cognitive performance by applying the coefficients of determination (*R*^*2*^) calculated for all possible pairs of variables in four groups of participants. In order to characterize the segregation and integration properties of the individual networks of CMV-positive or -negative men and women, we analyzed such network topology measures as clustering coefficient, characteristic path length, local and global efficiency (25, 26). With the aim of statistically comparing the network topology dynamics and to identifying the networks with optimal features of segregation and integration, we applied a rewiring procedure. To the best of our knowledge, simultaneous network analyses of multiple inflammation-related peripheral biomarkers and cognitive performance of older Cytomegalovirus-seropositive and -seronegative men and women have not been previously accomplished.

## Methods

### Participants

The sample has already been described in (22). It consisted of 161 older adults (Fig. 1) who had enrolled in a training study that included physical, cognitive, and combined training interventions. Male and female subjects were recruited from volunteer participant pools at the Max Planck Institute for Human Development and by advertisements in the metropolitan area of Berlin, Germany. All the volunteers lived independently at home, leading an active life. Participants were healthy, right-handed adults, aged 64-79 years. All volunteers completed a medical assessment prior to data collection. The medical examination was conducted at the Charité Sports Medicine, Charité Universitätsmedizin Berlin. Of the originally recruited 201 volunteers only 179 individuals met inclusion criteria for study participation after medical assessment. None of the participants had a history of head injuries, medical (e.g., heart attack), neurological (e.g., epilepsy), or psychiatric (e.g., depression) disorders. None of the volunteers had suffered from chronic inflammatory, autoimmune or cancer diseases, nor had clinically evident infections. Moderately elevated and controlled blood pressure was not considered as an exclusion criterion. All subjects completed the informed consent form to the study protocol which was approved by the Ethics Committee of the German Society of Psychology, UL 072014.

**Figure 1.**
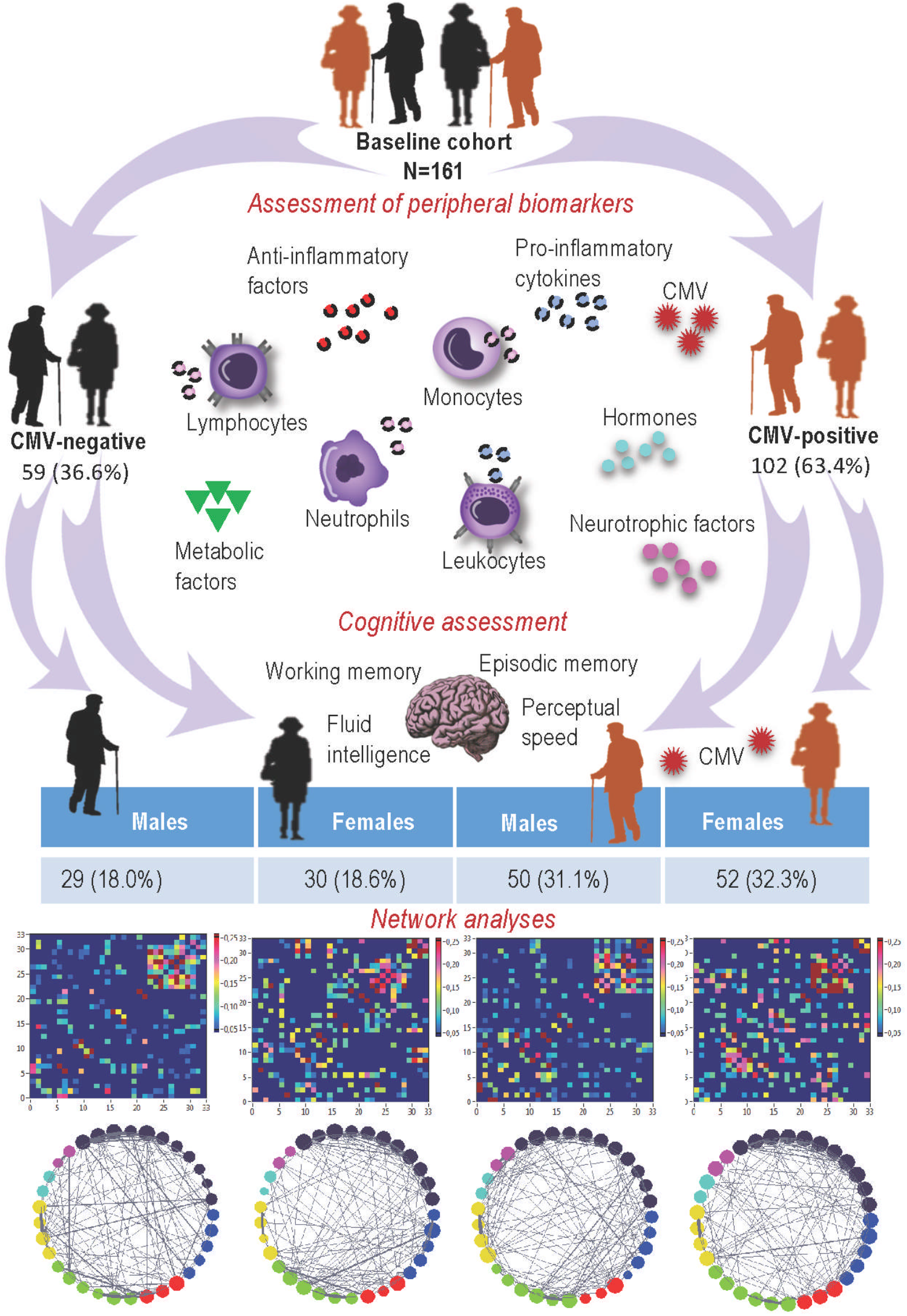
A schematic illustration of the study setup. Modified from (22). CMV: Cytomegalovirus.

### Circulating biomarkers assessment

The assessment of circulating cytokines, receptor antagonist, soluble cytokine receptor, and CMV-serostatus has been described in detail (22). For all analyses, the participants were separated into four groups according to their CMV-serostatus and sex (Fig. 1).

### Cytokines TNF, IL-10, IL-6, and IL-1β

Serum levels of pro- and anti-inflammatory cytokines (TNF, IL-10, IL-6, and IL-1β) were determined using the high-sensitivity cytometric bead array (CBA) flex system (BD Biosciences, San Jose, CA, USA) that allows multiplex quantification in a single sample. All analyses were performed according to the manufacturer’s instructions; to increase accuracy, an additional standard dilution was added. The fluorescence produced by CBA beads was measured on a BD FACS CANTO II Flow Cytometer and analyzed using the software FCAP Array v3 (BD Biosciences).

### sTNF-R, IL-1RA, IL-18, Cortisol, and DHEA levels, and CMV-serostatus

To gauge sTNF-R (80 kDA), IL-1RA, and IL-18 levels, we used the Sandwich Enzyme-linked Immunosorbent Assay (ELISA), a sensitive method allowing for the measurement of an antigen concentration in an unknown sample. All analyses were conducted according to the manufacturer’s instructions. The levels of human circulating sTNF-R (80 kDA), IL-1RA, and IL-18 were determined using the Platinum ELISA kit for the quantitative detection of the three cytokines (ThermoFisher SCIENTIFIC Invitrogen, Vienna, Austria, catalog numbers: BMS211, BMS2080 and BMS267/2).

Serum levels of anti-Cytomegalovirus IgG were determined using a commercial ELISA kit (IBL International GMBH, Hamburg, Germany, catalogue number: RE57061) and according to the manufacturer’s instructions. Samples were considered to give a positive signal if the absorbance value exceeded 10% over the cut-off, whereas a negative signal was absorbance lower than 10% below the cut-off.

Quantitative determination of Cortisol and DHEA in serum of participants was performed using Human Cortisol and Human DHEA (sulfate form) ELISA kits (Qarigo Biolabatories, catalog number: ARG81162 and ARG80837). The central mechanism of the competitive ELISA is a competitive binding process performed by sample antigen and add-in antigen. The amount of bound add-in antigen is inversely proportional to the concentration of the sample antigen. The analyses were performed according to the manufacturer’s instructions.

All samples were assessed in duplicate at 450 or 450/620 nm using a Multiscan-FC Microtiter Plate Photometer. Protein concentrations were determined in relation to a four-parameter standard curve (Prism 8 GraphPad, San Diego, CA, USA) or calculated using Microsoft Excel 2011.

### Levels of IGF-1 and IGFBP-3, CRP, metabolic factors, and immune cells

Serum levels of Insulin-like growth factor 1 (IGF-1) and Insulin-Like Growth Factor-Binding Protein 3 (IGFBP-3) were determined at the Institute of Clinical Neurobiology at the University of Würzburg. The analyses were performed according to the manufacturer’s instruction using the Immulite 2000 systems: IGF-1 (L2KIGF2); IGFBP-3 (L2KGB2), Siemens Healthcare, Germany - an automated solid-phase, Electrochemiluminescence-Immunoassay (ECLIA).

Levels of C-reactive protein (CRP), cholesterol, LDL, HDL, triglyceride, lymphocytes, leukocytes, monocytes, and neutrophils were measured within the clinical diagnostics facility of Berlin, Labor28. Serum concentrations of cholesterols and triglyceride were measured using enzymatic colorimetric tests (Roche, Basel, Switzerland). Counts of the immune cells were determined by flow cytometry (Sysmex, Norderstedt, Germany).

### Cognitive assessment

Participants were invited to a baseline session that lasted about 3.5 h, in which they were tested in groups of four to six individuals. The cognitive battery included a broad range of measures of learning and memory performance, processing speed, working memory, and executive functioning. The group received a standardized session protocol and started, after instructions, each task with practice trials to ensure that all participants understood the task. Responses were collected via button boxes, the computer mouse, or the keyboard. A detailed description of the tasks and scores used in the present study is included in the supplementary material.

### Network construction and network properties

For network construction, we used a coefficient of determination (*R*^*2*^), ranging between 0 and 1, and indicating the extent to which one dependent variable is explained by the other. The coefficient of determination was calculated between all pairs of variables (N = 33) for the four experimental groups separately. Thus, the common network in each of the groups contained 33 nodes altogether, covering all possible interactions between the variables or nodes. To be able to construct sparse networks with relatively stable network topology, we first investigated ordered (lattice) and random networks containing the same number of nodes and edges as the real networks. To do so, we randomized the edges in the real network to achieve a random network. As for the lattice network, we redistributed the edges such that they were laying close to the main diagonal and in the corner opposite to the main diagonal with increasing order of their weights. The lattice network reconstructed in such a way has the same number of nodes and edges as the initial real network but is characterized by ring or lattice topology incorporating nearest-neighbor connectivity (27). Random networks were constructed 100 times, and the network topology measures determined each time were averaged for further analyses. To investigate the network topology of the real networks in topology space between regular and random networks with different wiring cost levels, we constructed real and control (i.e., lattice and random) networks in the range of costs between 10 and 60% with a step of 1% of wiring costs (the ratio of the number of actual connections to the maximum possible number of connections in the network). We then decided to set the cost level to 25%, which resulted in sparse and at the same time stable network topology (see below).

### Degrees and strengths

The degree of a node provides information about the number of links connected to that node, and the strength reflects the overall strength of that node’s connections or weights. Thus, the strength could be considered as a weighted degree. Degree or strength of a node indicates the activity of that node, whereas the sum or mean of all degrees (strengths) represents the overall activity of the network. As *R*^*2*^ is a weighted symmetric measure, we obtained the node’s strength 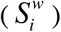 as the sum of weights of all connections (*w*_*ij*_) to node *i*, and calculated the mean strength (*S*) across all nodes in the network:

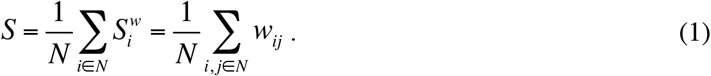

### Clustering coefficient and characteristic path length

For an individual node *i*, the clustering coefficient 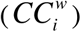 is defined as the proportion of the number of existing neighbor–neighbor connections to the total number of possible connections within its neighborhood. In the case of a weighted graph, the mean *CC* is calculated as follows (28):

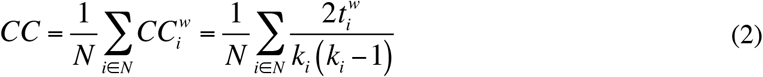

with 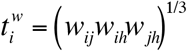 being the number of weighted closed triangles around a node *i*; *k*_*i*_ is the degree of the node *i*, and *N* is the number of nodes in the network, *N* =33. The *CC* measures the cliquishness of a typical neighborhood and is thus a measure of network segregation.

The shortest path length or distance *d*_*ij*_ between two nodes *i* and *j* is normally defined as the minimal number of edges that have to be passed to go from *i* to *j*. As our networks are weighted graphs, the weight of the links must be considered. The input matrix is then a mapping from weight to length (i.e., a weight inversion), and the distance 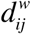 is the minimal weighted distance between the nodes *i* and *j*, but not necessarily the minimal number of edges. To calculate the characteristic path length (*CPL*) of a network, path lengths between all possible pairs of vertices or nodes in the network were determined (29) and then averaged among nodes:

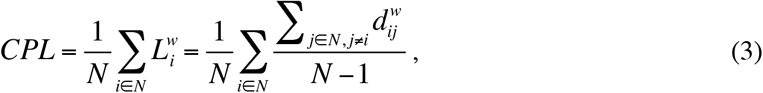

whereby 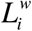 is the shortest path length of a node *i*, and *N* is the total number of nodes in the network. *CPL* shows the degree of network integration, with a short *CPL* indicating higher network integration.

### Local and global efficiency

Local efficiency (*E*_*local*_) is similar to the *CC* and is calculated as the harmonic mean of neighbor-neighbor distances (30):

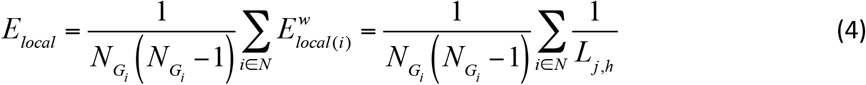

where 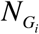 is the number of nodes in subgraph *G*_*i*_, comprising all nodes that are immediate neighbours of the node *i* (excluding the node *i* itself), and 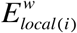 is local efficiency of the node *i* determined as the reciprocal of the shortest path length between neighbours *j* and *h*. Thus, *E*_*local*_ of node *i* is defined with respect to the subgraph comprising all of *i*’s neighbours, after removal of node *i* and its incident edges (Latora and Marchiori, 2001). Like *CC*, *E*_*local*_ is a measure of the segregation of a network, indicating efficiency of information transfer in the immediate neighbourhood of each node.

Global efficiency (*E*_*global*_) is defined as the average inverse shortest path length and is calculated by the formula (30):

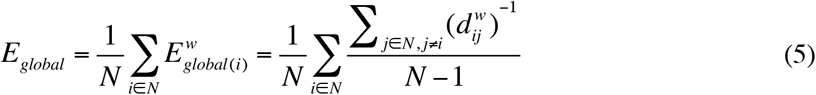

whereby 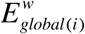 is a nodal efficiency, 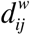 is the minimal weighted distance between the nodes *i* and *j*, and *N* is the total number of nodes in the network. The nodal efficiency is practically the normalized sum of the reciprocal of the shortest path lengths or distances from a given node to all other nodes in the network. Nodal efficiency quantifies how well a given node is integrated within the network, and global efficiency indicates how integrated is the common network. Thus, like *CPL, E*_*global*_ is a measure of the integration of a network, but whereas *CPL* is primarily influenced by long paths, *E*_*global*_ is primarily influenced by short ones.

### Small-Worldness (SW) Coefficients

Using graph metrics determined for real and control (i.e., regular and random) networks, specific quantitative small-world metrics were obtained. The first small-world metric, the so-called small-world coefficient σ, is related to the main metrics of a random graph (*CC*_*rand*_ and *CPL*_*rand*_) and is determined on the basis of two ratios γ = *CC*_*real*_ / *CC*_*rand*_ and *λ* = *CPL*_*real*_ / *CPL*_*rand*_ (31):

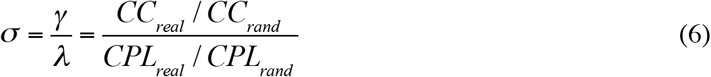

The small-world coefficient σ should be greater than 1 in the small-world networks (SWNs). The second SW metric, the so-called small-world coefficient *ω*, is defined by comparing the characteristic path length of the observed (real) and random networks, and comparing the clustering coefficient of the observed or real network to that of an equivalent lattice (regular) network (32):

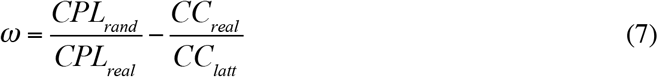

This metric ranges between −1 and +1 and is close to zero for SWN (*CPL*_real_≈*CPL*_rand_ and *CC*_real_≈*CC*_latt_). Thereby, negative values indicate a graph with more regular properties (*CPL*_real_>>*CPL*_rand_ and *CC*_real_≈*CC*_latt_), and positive values of ω indicate a graph with more random properties (*CPL*_real_≈*CPL*_rand_ and *CC*_real_<<*CC*_*l*att_). As suggested in (32), the metric ω compared to σ has a clear advantage, i.e., the possibility to define how much the network of interest resembles its regular or random equivalents.

### Modularity Analyses and Z-P Parameter Space

To investigate the modular organization of the network and the individual role of each node in the emerging modularity or community structure, we partitioned the networks into modules applying modularity optimization algorithm and determined indices of modularity (*Q*), within-module degree (*Z*_*i*_), and participation coefficient (*P*_*i*_) using the Brain Connectivity Toolbox (33). The optimal community structure is a subdivision of the network into non-overlapping groups of nodes in a way that maximizes the number of within-module edges, and minimizes the number of between-module edges. *Q* is a statistic that quantifies the degree to which the network may be subdivided into such clearly delineated groups or modules. It is given for weighted networks by the formula (34):

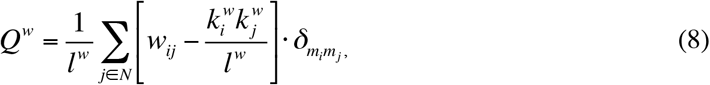

where *l*^*w*^ is the total number of edges in the network, N is the total number of nodes in the network, *w*_*ij*_ are connection weights, 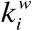 and 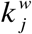 are weighted degrees or strengths of the nodes, and 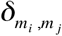 is the Kronecker delta, where 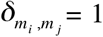 if *m*_*i*_ = *m*_*j*_, and 0 otherwise. High modularity values indicate strong separation of the nodes into modules. *Q*^*w*^ is zero if nodes are placed at random into modules or if all nodes are in the same cluster. To test the modularity of the empirically observed networks, we compared them to the modularity distribution (*N* = 100) of random networks as described above (35). The within-module degree *Z*_*i*_ indicates how well node *i* is connected to other nodes within the module *m*_*i*_. As shown in Guimerà and Amaral (36), it is determined by:

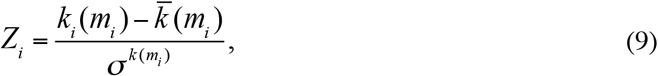

where *k*_*i*_(*m*_*i*_) is the within-module degree of node *i* (the number of links between *i* and all other nodes in *m*_*i*_), and 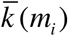 and 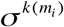 are the mean and standard deviation of the within-module degree distribution of *m*_*i*_. The participation coefficient *P*_*i*_ describes how well the nodal connections are distributed across different modules (36):

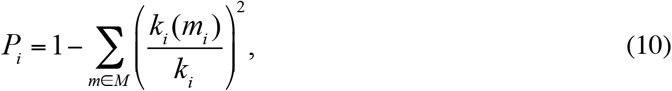

where *M* is the set of modules, *k*_*i*_*(m*_*i*_) is the number of links between node *i* and all other nodes in module *m*_*i*_, and *k*_*i*_ is the total degree of node *i* in the network. Correspondingly, *P*_*i*_ of a node *i* is close to 1 if its links are uniformly distributed among all the modules, and is zero if all of its links lie within its own module. *Z*_*i*_ and *P*_*i*_ values form a so-called *Z-P* parameter space and are characteristic for the different roles of the nodes in the network (36). These roles in the *Z-P* parameter space could be defined as follows: ultra-peripheral nodes (*P*_*i*_<0.05), provincial nodes (low *Z*_*i*_ and *P*_*i*_ values), connector nodes (low *Z*_*i*_ and high *P*_*i*_ values), hub nodes (high *Z*_*i*_ and low *P*_*i*_ values), and hub connector nodes (high *Z*_*i*_ and *P*_*i*_ values). In this context, hubs are responsible for intra-modular connectivity and contain multiple connections within a module, while connector nodes maintain inter-modular connectivity and are responsible for links between the modules.

### Statistical analysis

In order to statistically compare the four different networks at a given cost level, we used a rewiring procedure with a step-by-step replacement of a non-existing edges through an existing ones and consecutive determination network topology metrics each time. This procedure can specify the network stability and network topology alteration by very small changes in the network configuration. In a statistical sense, this procedure is similar to bootstrapping with replacement applied to time series. In total, there were about 50,000 rewired networks, on which mean and standard deviation (SD) of the network topology metrics were determined. Because the rewiring distribution showed a normal shape and a small bias, we were able to achieve a 99.7% confidence interval (*CI*) for the mean by using the empirical rule: *CI* = *mean* ± *3* × *SD* (*P* < 0.005).

## Results

### Network composition and network topologies in real and control networks

Before analyzing network topology changes, we compared the topology in real and control (i.e., lattice and random) networks under different cost levels (i.e., in the range of wiring costs between 10 and 60%). As shown in Supplementary Figure 1A, *CC* is greatest in lattice networks and lowest in random networks, whereas *CC* for the real networks lies in-between. *CPL* is shortest in random and longest in lattice networks, while the real networks are between these (see Supplementary Fig. 1B). Correspondingly, *E*_*local*_ was highest in lattice networks (at least for cost levels under 45%) and lowest in random networks (at least for cost levels under 20%), while *E*_*global*_ was highest in random and lowest in lattice networks essentially for all levels of wiring costs, with real networks always in between (see Supplementary Fig. 2 for details).

Importantly, as shown in Figure 2, networks under consideration are Small-Word Networks (SWNs) at all levels of wiring costs (σ > 1). As indicated by the other SW coefficient ω, which is lying at practically all levels of wiring costs in the positive range (see Fig. 2B), these networks are SWNs with more random characteristics. It can also be seen that the networks with costs lower than 25% showed rather unstable behavior that was stabilizing at the 25% level of costs and showed very similar results across all experimental groups for both SW coefficients σ and ω. Thus, for our main analyses, we decided to set the cost level to 25% that makes it possible to investigate sparse and at the same time stable network topology in all four groups of participants.

**Figure 2.**
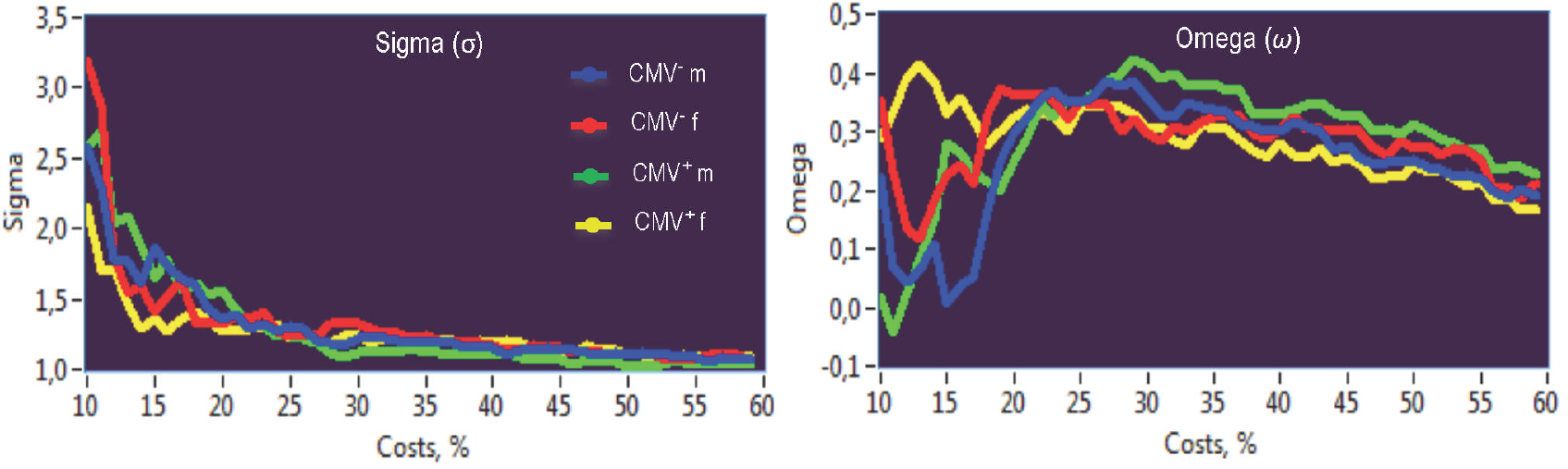
Small-world coefficients sigma (σ) and omega (*ω*) under different levels of the wiring costs. CMV, Cytomegalovirus; CMV^−^ m, CMV-seronegative men; CMV^+^ m, CMV-seropositive men; CMV^−^ f, CMV-seronegative women; CMV^+^ f, CMV-seropositive women.

### Network structure and network strengths

It can be seen that connectivity matrices (Fig. 3A) display a group-specific structure in all four participant groups. In the first step, we calculated network strengths as the sum of connections of node *i*.

**Figure 3.**
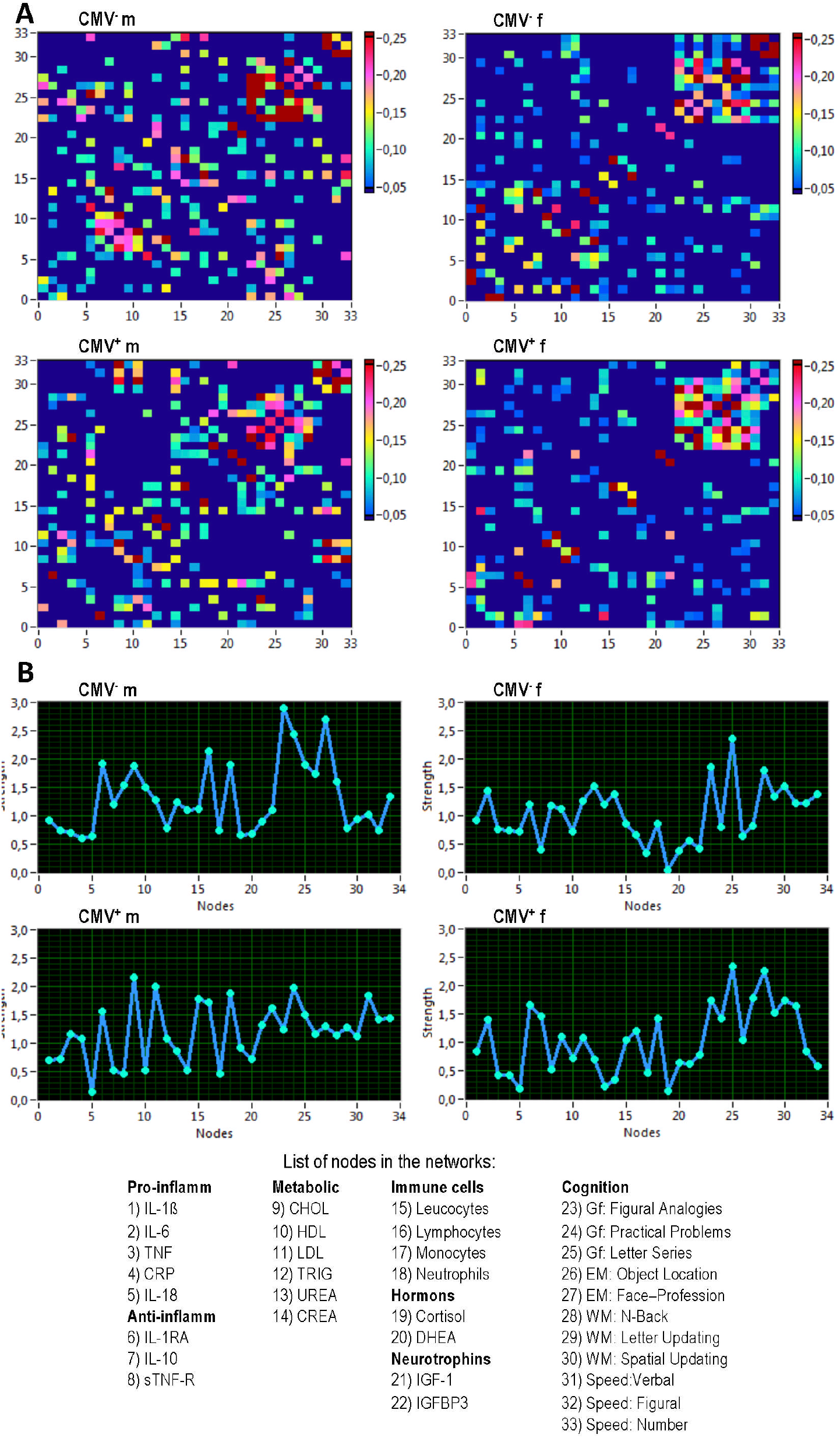
Connectivity structure of the network and network strengths in the four groups. (**A**) Connectivity matrices. (**B**) Network strengths. CMV, Cytomegalovirus; CMV^−^ m, CMV-seronegative men; CMV^+^ m, CMV-seropositive men; CMV^−^ f, CMV-seronegative women; CMV^+^ f, CMV-seropositive women; IL, interleukin; IL-1β, interleukin 1 beta; TNF, tumor necrosis factor; CRP, C-reactive protein; IL-1RA, interleukin 1 receptor antagonist; sTNF-R, soluble tumor necrosis factor receptor; CHOL, cholesterol; HDL, high-density lipoprotein; LDL, low-density lipoprotein; TRIG, triglyceride; CREA, creatinine; DHEA, dehydroepiandrosterone; IGF-1, insulin-like growth factor-1; IGFBP-3, IGF-binding protein 3; Gf, fluid intelligence; EM, episodic memory; WM, working memory; Speed, perceptual speed.

As shown in Figure 3 (A-B), cognitive nodes exhibit high strengths, which are mostly due to the strong connections between the cognitive nodes themselves, especially in the female groups. In the male groups, the cognitive nodes are also strongly connected to the other systems such as cytokines (especially, in the network of CMV^−^ males), metabolic variables (particularly, in the network of the CMV^+^ males) and immune cells.

### Networks of CMV- and CMV+ men and women differ in their structure

Networks of the four experimental groups also display group-specific structure (Fig. 4). Individual nodes (or variables) are represented as multicolored circles coding for affinity to a particular group of variables.

**Figure 4.**
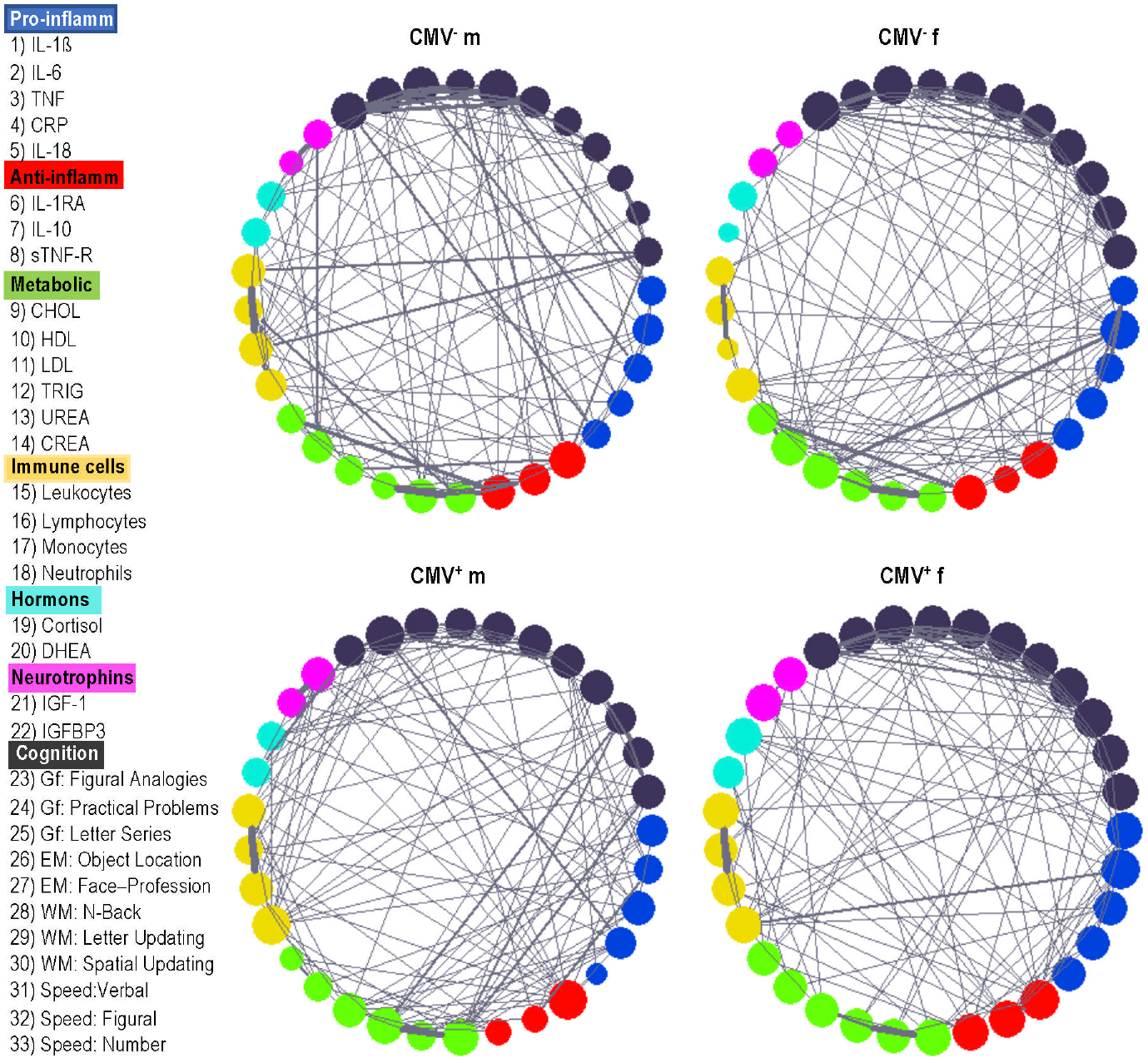
Network structure differences in CMV^−^ and CMV^+^ men and women. CMV, Cytomegalovirus; CMV^−^ m, CMV-seronegative men; CMV^+^ m, CMV-seropositive men; CMV^−^ f, CMV-seronegative women; CMV^+^ f, CMV-seropositive women; IL, interleukin; IL-1β, interleukin 1 beta; TNF, tumor necrosis factor; CRP, C-reactive protein; IL-1RA, interleukin 1 receptor antagonist; sTNF-R, soluble tumor necrosis factor receptor; CHOL, cholesterol; HDL, high-density lipoprotein; LDL, low-density lipoprotein; TRIG, triglyceride; CREA, creatinine; DHEA, dehydroepiandrosterone; IGF-1, insulin-like growth factor-1; IGFBP-3, IGF-binding protein 3; Gf, fluid intelligence; EM, episodic memory; WM, working memory; Speed, perceptual speed.

The size of the circle depends on the sum of connections and indicates the node’s strength. The thickness of the connections corresponds to their connection strength. The nodes are numbered clock-wise beginning from the pro-inflammatory cytokine IL-1β displayed in blue. The CMV-negative male group (top, left) is characterized by multiple strong connections between pro-inflammatory cytokine nodes (IL-1β, TNF, IL-18) and cognitive nodes (episodic memory and fluid intelligence).

Less strong but numerous connections are also present for anti-inflammatory cytokines and the cognitive nodes. Interestingly, this is the only group, in which pro- and anti-inflammatory cytokines show no direct connections to each other. The nodes of perceptual speed are strongly connected with immune cell nodes (lymphocytes and neutrophils). No other groups of participants display such strong direct connections between immune biomarkers and cognition – except the network of CMV^+^ men (bottom, left) with only one strong connection between CRP and fluid intelligence. The network of the CMV^+^ men shows strong connections between metabolic factors and perceptual speed. The network of CMV^−^ women (top, right) displays strong connections between pro-inflammatory IL-6 and triglycerides as well as between anti-inflammatory sTNF-R and creatinine. The network of the CMV^+^ women (bottom right) shows a strong connection between leukocytes and pro-inflammatory IL-6. Unexpectedly, neurotrophins in the CMV^−^ men have relatively strong connections to urea, but only one weak connection to the pro-inflammatory factor CRP. In contrast, all three of the other networks display multiple connections to both pro- and anti-inflammatory cytokines. Concerning connections between neurotrophins and cognitive nodes, we can see quite heterogeneous picture: with some connections in CMV-seronegative and -positive men, and with only one connection in the CMV-seronegative and - positive women. In general, the networks of all groups of participants show strong (but differently manifested) connections between the cognitive nodes themselves (Fig. 4).

### Networks topology differences between CMV^−^ and CMV^+^ men and women

To be able to statistically compare the four different networks at a given cost level, we used rewiring procedure with replacement of a non-existing edge through an existing one and consecutive determination of network topology metrics each time. In total, there were about 50,000 rewired networks, for which mean and standard deviation (SD) of the network topology metrics were determined. In accordance with the empirical rule, we achieved a 99.7% confidence interval (*CI*) for the mean: *CI* = *mean* ± *3* × *SD*. As shown in Figure 5 (A), mean *CC* was highest and *CPL* shortest in CMV^−^ males and in total, higher (shorter) in males than in females. Correspondingly, local and global efficiency were both highest in CMV^−^ males and in total higher in males than in females. CMV-seronegative and - seropositive females did not show any significant differences. This indicates that segregation and integration properties of the network were notably stronger in males (especially, in CMV^−^ males) than in females. Inspection of separate nodes in the networks showed that these network topology differences were in particular stronger for cytokines and cognitive variables or nodes (Figure 5, B).

**Figure 5.**
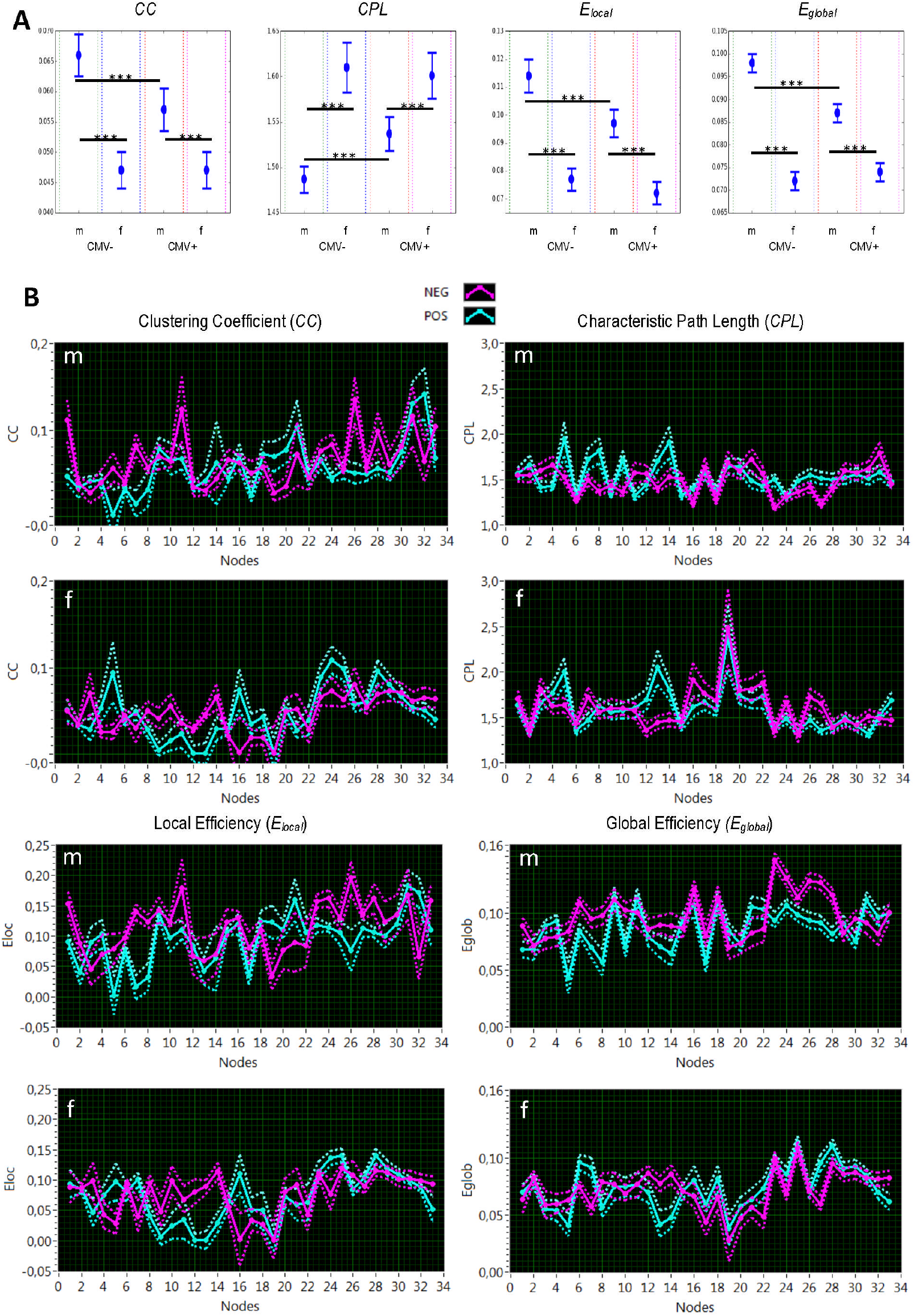
Network topology differences. (**A**) Results of rewiring analyses for whole network. (**B**) Results of rewiring analyses for individual nodes. *CC*, clustering coefficient; *CPL*, characteristic path length; *E*_*local*_, local efficiency; *E*_*global*_, global efficiency; CMV, Cytomegalovirus; CMV-, CMV-seronegative; CMV+, CMV-seropositive; m, male; f, female; NEG, CMV-seronegative; POS, CMV-seropositive.

### Modular organization of the networks of CMV^−^ and CMV^+^ men and women

Modularity analyses showed that the networks under consideration exhibited in all cases highly differentiated modular organization with 4 and 5 modules for males and for females, respectively. This is indicated by high modularity values or Q statistics (Fig. 6), which ranged between 0.397 and 0.453, and were considerably higher as compared with random networks (with *Q*-values close to 0). Nodes sharing the same module are displayed in Figure 6 (B and D) in the same color.

**Figure 6.**
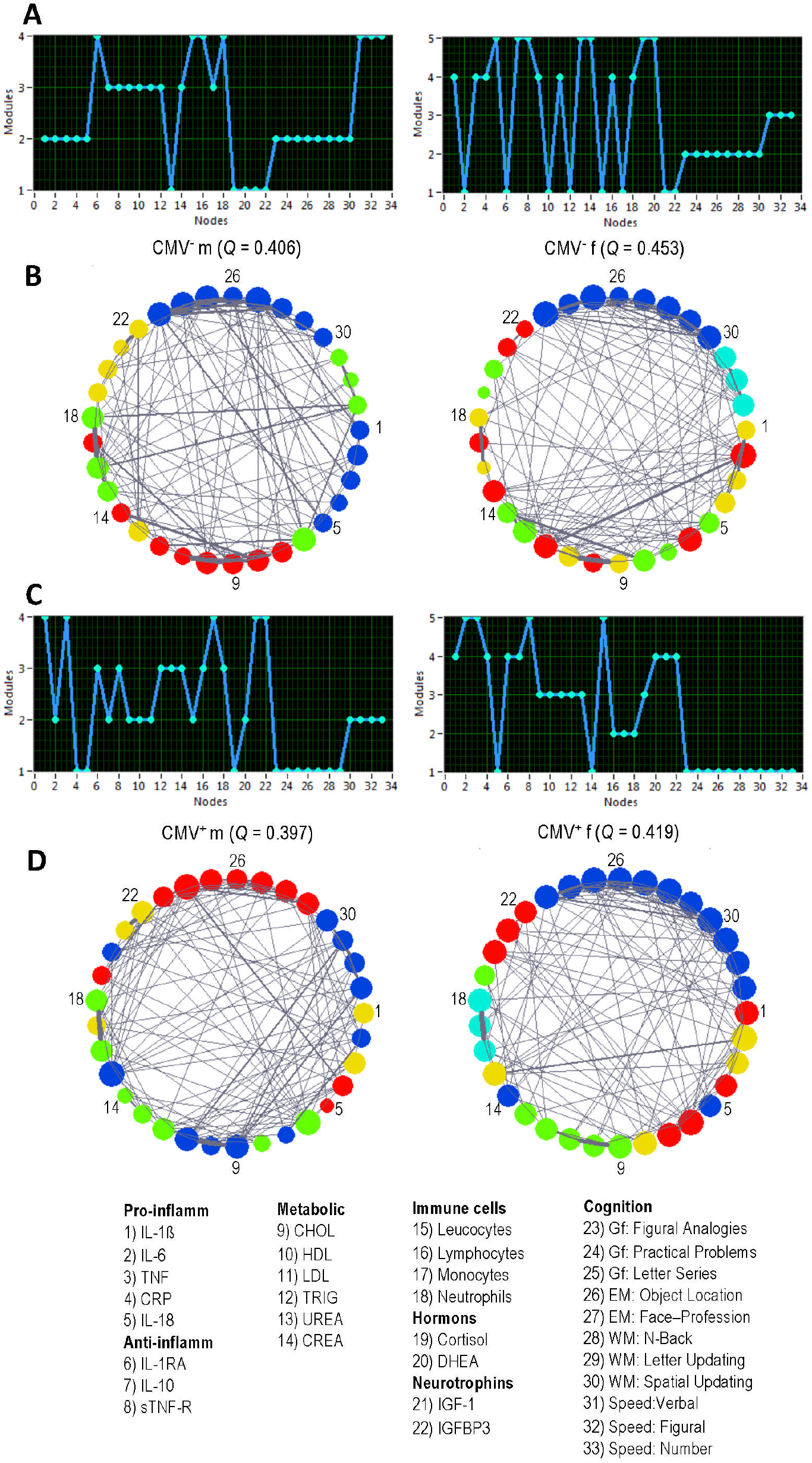
Modular organization of the networks. (**A**) Modular assignment of nodes in CMV^−^ men (left) and women (right). (**B**) Modular structure in CMV^−^ men (left) and women (right). (**C**) Modular assignment of nodes in CMV^+^ men (left) and women (right). (**D**) Modular structure in CMV^+^ men (left) and women (right). Note that nodes sharing the same module are displayed in the same color. CMV, Cytomegalovirus; CMV^−^ m, CMV-seronegative men; CMV^+^ m, CMV-seropositive men; CMV^−^ f, CMV-seronegative women; CMV^+^ f, CMV-seropositive women; Q, modularity value.

As shown in Figure 6 (A and C), cognitive nodes occupied two modules in all networks (with exception of CMV^+^ females, in which all cognitive nodes were located in one large module), whereby perceptual speed nodes occupied a separate module. Moreover, the community structure in CMV-negative males was organized in 4 modules (A-B, left), whereby all pro-inflammatory cytokines were located in the same module shared (B, blue) with cognitive variables or nodes (reflecting general intelligence and memory features). In addition, two of the three anti-inflammatory cytokines (namely, IL-10 and sTNF-R) shared the same module (B, left, red) with metabolic factors as well as with monocytes, with the exception of urea, which was located in a separate module (B, yellow) together with hormones und neurotrophins. Finally, perceptual speed nodes shared a common module (B, left, green) with IL-1RA and immune cells (namely, leukocytes, lymphocytes, and neutrophils). Interestingly, in CMV^−^ females (A-B, right), the two modules occupied by cognitive (B, right, blue) and perceptual speed nodes (B, right, cyan) were separated from all the other nodes, which were partitioned into heterogeneous modules comprising different components (e.g., cytokines, metabolic variables, immune cells, and neurotrophins). The nodes of CMV^+^ men (C-D, left) and CMV^+^ women (C-D, right) also partitioned into 4 and 5 modules, respectively, showed heterogeneous modularity structures comprising nodes of both peripheral biomarkers and cognitive features.

### *Z-P* Parameter space and nodes’ specificity of the four networks

To define how the network nodes were positioned in their own module and with respect to other modules, we calculated the within-module degree (*Z*_*i*i_) and participation coefficient (*P*_*i*i_) of the node *i* for the given networks. The within-module degree indicates how ‘well-connected’ node *i* is to other nodes in the module, whereas the participation coefficient reflects how ‘well-distributed’ the edges of the node *i* are among the other modules. *Z*_*i*_ and *P*_*i*_ form together the so-called *Z-P* parameter space, with different regions indicating specific roles of the nodes (e.g., hubs, connectors, provincial nodes) in this parameter space (36). As shown in Figure 7 (A), the network of the CMV^−^ males contains more hub nodes but far fewer connector nodes than the other three groups. This indicates that the modules in this participants’ group are more autonomous and the information flow between the modules is either reduced or is realized through a small number of connector nodes. Interestingly, three of the four hubs are cognitive variables and the fourth one is IGFBP3. Thus, cognitive nodes, such as fluid intelligence, working memory, and perceptual speed, play a central role in the network of CMV^−^ males driving or controlling the connections within the corresponding modules. Further, the networks of CMV^−^ females (B) and CMV^+^ males (C) are characterized by high numbers of the non-hub connectors responsible for the connectivity between the modules.

**Figure 7.**
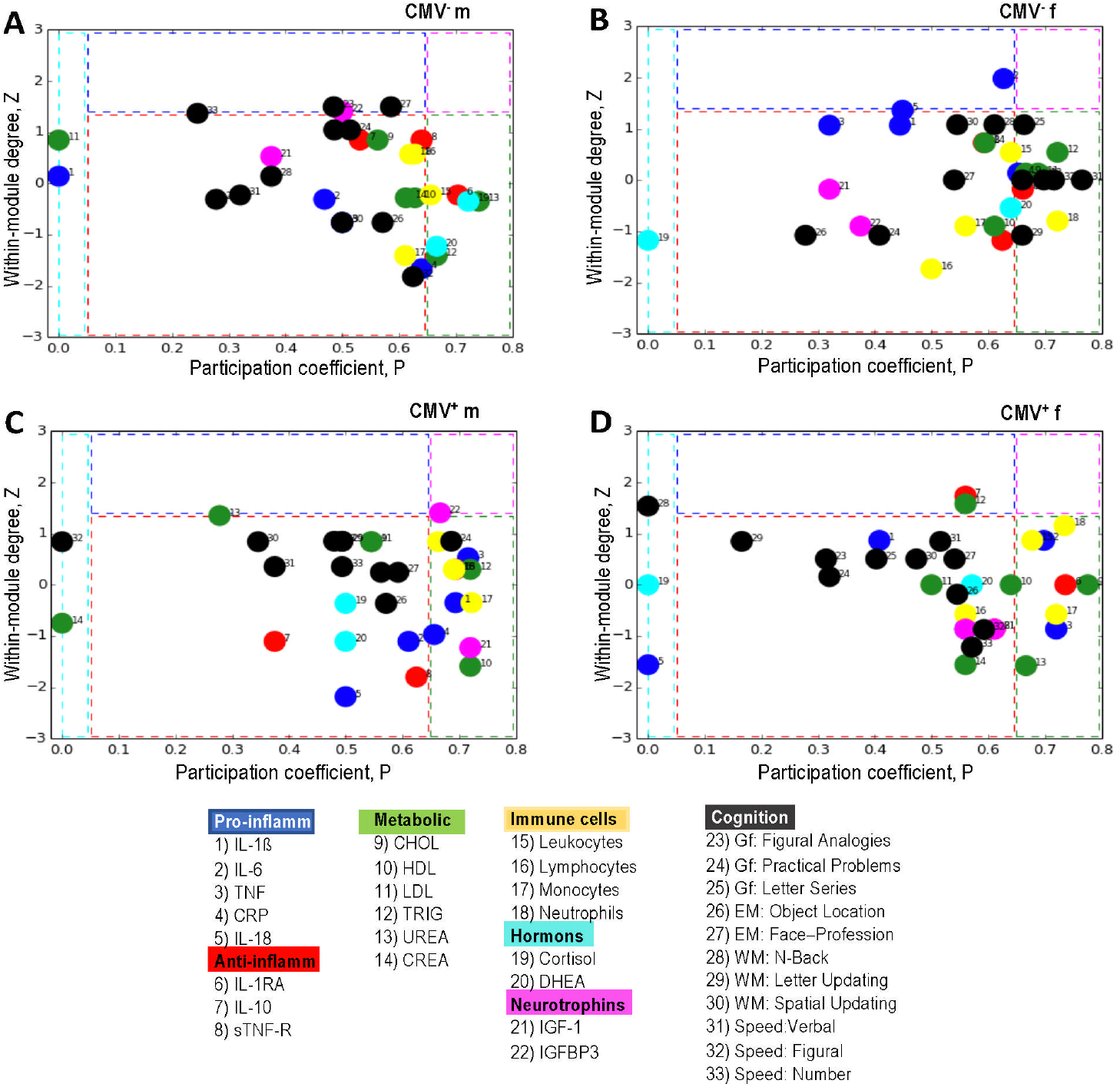
*Z-P* parameter space and node’ specificity for networks in four groups. (**A**) *Z-P* parameter space for CMV-seronegative men, (**B**) *Z-P* parameter space for CMV-seronegative women, (**C**) *Z-P* parameter space for CMV-seropositive men, and (**D**) *Z-P* parameter space for CMV-seropositive women. Different regions separated by dotted lines contain: left – ultra-peripheral nodes; central – provincial nodes; top – hubs; top right – connector hubs; right – connectors. CMV, Cytomegalovirus; CMV^−^ m, CMV-seronegative men; CMV^+^ m, CMV-seropositive men; CMV^−^ f, CMV-seronegative women; CMV^+^ f, CMV-seropositive women.

Thus, the modules in these two groups are apparently worse separated from each other than, for example, in the CMV^−^ males. The network of the CMV^+^ females (D) contains two hubs and eight non-hub connectors, and thus demonstrates a modular structure with moderate number of hubs and connectors. Note also that all cognitive nodes in this group are provincial nodes and therefore play a secondary role in the network. In summary, it can be stated that the networks under consideration exhibit a different balance between intra- and inter-modular information flow with different numbers of hub and connector nodes playing a significant role for this balance and for network functioning. Which of these types of modular organization is more effective, remains to be investigated.

## Discussion

There is a growing body of evidence supporting the notion that the immune system is not hermetically self-regulated but functions in intimate interrelations with other physiological systems, including the nervous system (5, 37). These interactions are present at the various levels of organization – at the local, as well as at the whole organism level – by sharing a common language of a wide range of cytokines, receptor molecules, hormones, neuropeptides, metabolic and neurotrophic factors allowing cross-communication (38, 39). Particularly in the process of aging, this reciprocal cross-talk may under certain circumstances permit augmentation of maladaptive inflammatory loops, which could disturb homeostasis and contribute to the age-related functional alterations or even to pathological conditions (2, 40, 41).

Several analytical techniques to investigate these interactions have been established so far, but our understanding of the interplay between different factors in such interrelated processes is still in its infancy. Despite some progress, there is a further need to place the data from different physiological and functional levels in a biological context with the aim of interpreting their multifaceted orchestration as a whole. Many studies highlight the role of different inflammatory cytokines in the low-grade inflammation, dubbed “inflammaging”, and the importance of pro-inflammatory and anti-inflammatory homeostasis for cognitive health in aging (17, 18, 42-44). Additionally, the interrelated effects of inflammatory factors and their influence on neuroimmune and neuroendocrine functions can be modified by the chronic immune activity required to control lifelong persistent CMV infection (2, 45). In the present work, we propose a strategy for quantitative description of multiple interactions between different cytokines, receptor molecules, metabolic and neurotrophic factors, hormones, immune cells, and measures of cognitive performance with the help of a graph-theoretical approach. To the best of our knowledge, simultaneous network analyses of multiple inflammation-related mediators and cognitive performance in older CMV-seropositive and CMV-seronegative men and women have not been previously accomplished.

The fact that CMV has considerable influence on immunosenescence was first described 20 years ago (46) and has continuously been supported by numerous studies since then (15, 16, 47-52). In the large-scale immune profiling and functional analysis of normal aging, it was impressively shown that the immune system alterations (determined as a number of significantly affected analytes) caused specifically by CMV, were comparable with the differences seen between the sexes (53). A lifelong persistent infection influences immune aging and can significantly modify the course of cognitive aging by acting in combination with individual differences in cytokine release (45, 54-56). The modulatory effect of CMV-latency and sex were also demonstrated in our previous study (22). Therefore, for the network analyses in the present study, we separated the participants into four groups according to their CMV-serostatus and sex.

We found that the modulatory impact of CMV and sex was also reflected in the specific differences of the network structure and the network topology dynamics observed between the four groups. In particular, CMV^−^ males were characterized through several strong connections between nodes of the pro-inflammatory cytokines IL-1 β, TNF, IL-18 and cognitive nodes including variables of episodic memory and fluid intelligence. Currently available evidence shows that pro-inflammatory cytokines exert a dose-dependent physiological neuroprotective but can however also mediate pathological neurodegenerative effects under certain circumstances (18). IL-1β and TNF were demonstrated to have such a dual function, acting on the one hand as pro-inflammatory factors and on the other as neuromodulators, subserving memory and other cognitive processes. In other words, they not only play a role in neuroinflammation, but (at their low concentrations) also in complex processes such as synaptic plasticity, neurogenesis, long-term potentiation and memory consolidation (42, 43).

Less strong but numerous connections were found between nodes of the anti-inflammatory cytokines and cognition in the network of CMV^−^ males. This is partly in line with our previous findings on the positive association of episodic memory with the anti-inflammatory cytokine IL-10 in the CMV^−^ elderly men and women (22). IL-10 is known to have a neuroprotective role due to its inhibitory action on inflamed microglia (17). The same CMV^−^ male group also has significantly elevated levels of anti-inflammatory IL-10 and sTNF-R as well as reduced levels of pro-inflammatory cytokines in their peripheral circulation, as reported in our recent study (22). Having this information in mind, we can speculate that strong connections between cognitive nodes and the nodes of (low-levels) pro-inflammatory cytokines on the one hand and numerous connections of cognition to the nodes of the (high-level) anti-inflammatory cytokines on the other, could possibly explain the cognitive advantage in the fluid intelligence and working memory found for this group of participants in our previous work (22). Remarkably, this was the only group in which nodes of pro- and anti-inflammatory cytokines had no direct connections to each other. The other three groups, (two of which, CMV^−^ females and CMV^+^ males, were characterized in our previous study by heterogeneously unbalanced levels of pro- and anti-inflammatory mediators and by an adverse metabolic environment) demonstrated, in contrast, various more or less strong connections between pro- and anti-inflammatory cytokines, which were probably important and necessary homeostatic responses to these unbalanced peripheral conditions. In our previous study, the network of CMV^+^ women (that shows multiple connections between nodes of pro- and anti-inflammatory cytokines), exhibited significantly higher levels of the anti-inflammatory factors sTNF-R and IL-1RA. We also found previously that in the CMV^+^ group, fluid intelligence, episodic and working memory were negatively associated with the anti-inflammatory factor IL-1RA, the level of which was assumed to be simultaneously increased as a reaction to the elevation of the pro-inflammatory cytokines in the periphery (22). This phenomenon has also been reported by other investigators (57, 58), showing that individuals with high levels of the pro-inflammatory cytokines also tend to display elevated levels of anti-inflammatory factors. The network analyses in the present study allowed the visualization of these multiple and mutual connections between pro- and anti-inflammatory biomarkers, which were only assumed in our previous work (22).

Interestingly, the network of CMV-males demonstrated some direct connections between DHEA and cognitive nodes, and also to the nodes of anti-inflammatory and metabolic factors. The CMV^+^ males, in contrast, displayed multiple connections to cognitive nodes, but no connections to anti-inflammatory nodes, and were connected to the inflammatory cytokine IL-6. A completely different picture was seen in CMV^−^ females with no connections of DHEA either to pro-inflammatory cytokines or cognition, whereas CMV^+^ women had multiple connections to nodes of cytokines and cognition. It is known that inflammatory reactions are, in general, under the influence of different mechanisms including neuroendocrine interactions. Pro-inflammatory mediators and cytokines may lead to the activation of the hypothalamic-pituitary-adrenal axis (HPA) that is in turn capable of modulating the process of inflammation (59–63). DHEA and cortisol are multifunctional adrenocortical hormones with such immunomodulatory properties. They exert potent and broad influences throughout the body and brain and jointly impact on a variety of processes related to metabolic, immune, and cognitive functions (60). Being especially abundant in the brain, DHEA exerts a protective effect against the deterioration of mental functioning with aging. Interestingly, both cortisol and DHEA in the CMV^−^ males are non-hub connectors exhibiting numerous links to diverse modules in the modular organization of the network. This indicates that these nodes play a crucial role in communication between different subsystems. Inverse correlations between DHEA concentrations and neuroinflammatory-related diseases have repeatedly been found in the elderly (60, 64-66). Similar to DHEA, the cortisol nodes in our study displayed very heterogeneous and group-specific picture concerning their connections. Whereas CMV^−^ males showed connections from cortisol to the nodes of pro-inflammatory TNF, IGF-1, IGFBP-3, metabolic factors, and immune cells, the cortisol-node of CMV^−^ females had only one connection to IL-18. In the CMV^+^ groups, men showed weak but multiple cortisol-connections to cognitive nodes, neurotrophins, pro- and anti-inflammatory factors. In the network of women, cortisol was connected only to the metabolic factors. The heterogeneous picture seen in these connections may partly be due to the fact that although the effect of cortisol has been typically shown to be immunosuppressive, at certain concentrations it can also induce a biphasic response during a later, delayed systemic inflammatory response (67) through augmentation of inflammation (61). In other words, the regulation of inflammation by cortisol may vary from anti-to pro-inflammatory in a time- and concentration-dependent manner and this contributes to further complexity in interpreting results of these already complex interactions.

Pro-inflammatory cytokines are known to be involved in dynamic interactions with the main neurotrophic factor, IGF-1 and its regulator, IGFBP-3 by decreasing IGF-1 signaling and by enhancing the production of IGFBP-3. Conversely, IGF-1 is capable of depressing pro-inflammatory cytokine signaling by increasing anti-inflammatory IL-10 secretion and by directly depressing pro-inflammatory cytokine signaling (23, 68, 69). Both IGF-1 and IGFBP-3 had relative strong connections to metabolic nodes in the CMV-men, but only one weak connection to CRP. In contrast, all three of the other networks displayed multiple connections to both pro- and anti-inflammatory cytokines – possibly due to their involvement in the dynamic interactions aiming to balance the pro- and anti-inflammatory equilibrium. Concerning the connections between neurotrophins and cognitive nodes, we can see a relative homogeneous picture: with some connections in the networks of CMV-negative and -positive men, and with only one connection in the networks of CMV-negative and -positive women. There is substantial evidence that IGF-1 deficiency represents a contributing factor for reduced cognitive abilities in aged humans (65, 70), and that supplementation with IGF-1 may reverse this deficit (68, 71-74). Measures of circulating IGF-1, IGFBP-3 and their ratio, have been proposed for monitoring aged individuals and those at risk of cognitive and functional decline (70). Thus, we can speculate that the relatively low number of connections between neurotrophins and cognitive nodes, seen in all four networks, might be due to the overall age-related decrease of these neurotrophic factors in peripheral circulation of elderly participants.

Our study has many strengths, including that it is one of the first studies to extensively characterize, prior to any physical, cognitive, and combine interventions, the network topology dynamics in multiple peripheral circulating biomarkers and markers of cognitive functioning. Applying a graph-theory approach allowed us not only to visualize biologically meaningful interconnections between nodes but also for the first time to compare the network topology metrics between different groups of CMV-seronegative and -positive men and women in a statistically sound manner. Inspection of separate nodes in the networks showed that these network topology differences were especially strong for cytokines and cognitive nodes. Modularity analyses showed that the networks under consideration exhibited highly differentiated modular organization in all cases. Moreover, we found that all four networks represented so-called small-world networks (SWNs) at all levels of wiring costs and were identified as SWNs with more random characteristics. We found that the network of the CMV-males contains more hub nodes but fewer connector nodes than the other three groups. This indicates that the modules in this participants’ group are more autonomous and the information flow between the modules may be realized through a small number of connector nodes. Interestingly, three of the four hubs are cognitive variables and the fourth one is IGFBP-3. Thus, cognitive nodes, such as fluid intelligence, working memory, and perceptual speed play a central role in the network of CMV-males driving or controlling the connections within the corresponding modules.

This is the first study investigating the segregation and integration properties of the individual networks of CMV-seropositive and -negative older men and women by analyzing such network topology measures as clustering coefficient, characteristic path length, local and global efficiency. Using the rewiring procedure for network analyses, we compared network topology dynamics and found that mean clustering coefficient was highest and CPL shortest in the network of the CMV-males. The same network also manifested the highest local and global efficiency, allowing it to be identified as the network with optimal features of segregation and integration. In our previous study, the same group of participants displayed the most balanced inflammatory status in their peripheral circulation (with low levels of pro-inflammatory cytokines and high levels of anti-inflammatory biomarkers) as well as significantly higher cognitive performance in working memory and fluid intelligence (22). Further studies, however, are required to confirm these findings and to better understand such complex relationships and network topology changes between different groups of older CMV-seropositive and - negative men and women.

There are several limitations to our study that should be acknowledged. The first one has already been mentioned in our previous publication and is “related to the fact that our pre-training cohort consisted of relatively healthy, non-obese, and well-educated Berlin residents with a comparatively low seroprevalence for CMV for this age. For this reason, the generalizability of some of our findings may be limited to the Berlin healthy aging population or to a similar European population in urban areas” (22). Another limitation is related to the exploratory character of our study of the network patterns and their relationships. We are well aware that our choice of variables in the present study, selected on the basis of their involvement in the known age-related functional alterations in the immune, nervous, and other central physiological systems, does not necessarily cover all potential players and, we therefore need further more extended network analyses to obtain a more comprehensive picture on their dynamic interactions.

## Conclusions

Network analyses applying a graph-theoretical approach provide a useful strategy for visualization and quantitative description of multiple interactions between various circulating pro- and anti-inflammatory biomarkers, hormones, neurotrophic and metabolic factors, immune cells, and measures of cognitive performance and can be in general applied for analyzing interactions between different physiological systems. Applying this approach, we were also able to confirm our previous findings that CMV-infection and sex modulate multiple circulating biomarkers and cognitive performance and that balanced inflammatory and metabolic status in elderly contributes to better cognitive performance. Analyzing the network topology dynamics of circulating biomarkers and cognitive performance in older CMV-seropositive and -seronegative men and women we were able to show that highly integrated and segregated networks have optimal neuroimmune and cognitive interactions.

## Abbreviations

CMV: Cytomegalovirus
CC: Clustering coefficient
CPL: Characteristic path length
*E*_*local*_: Local efficiency
*E*_*global*_: Global efficiency
DHEA: Dehydroepiandrosterone
IGF-1: Insulin-like growth factor-1
IGFBP-3: IGF-binding protein
IL: Interleukin
IL-1RA: Interleukin 1 receptor antagonist
TNF: Tumor Necrosis Factor
sTNF-R: Soluble Tumor Necrosis Factor receptor
CRP: C-reactive protein
HDL: High-density lipoprotein
LDL: Low-density lipoprotein
IgG: Immunoglobulin G
CI: Confidence interval
EM: Episodic memory
WM: Working memory
Gf: Fluid intelligence
IGF-1: Insulin-like growth factor 1
ELISA: Enzyme-linked Immunosorbent Assay
CBA: Cytometric bead array

## Acknowledgments

We would like to express our very great appreciation to Elisabeth Wenger for her valuable, constructive, and helpful suggestions during the study. We thank Sandra Düzel for providing cognitive data, reading manuscript, and her constructive remarks. We thank Marcel Gaetjen for his excellent methodological support in applying of the CBA-flex system and for providing the FCAP-Array-v3 software. We are thankful to the students of the Structural Plasticity Group for their great contribution in collecting the data reported above. We would like to thank Nadine Taube, Kirsten Becker, and Anke Schepers-Klingebiel for managing all organizational issues. We thank Carola Misgeld for medical data assessment and blood collection. We are grateful to all participants of the study.

## Availability of data and materials

The datasets for this study will not be made publicly available due to restrictions included in the consent statement that the participants of the study signed only allow the present data to be used for the research purposes within the Max Planck Institute for Human Development in Berlin.

## Author contributions

Conceptualization: S.D.B., L.M., G.P., M.S., and V.M.; methodology: V.M., S.D.B.; S.R., and M.F.; software: V.M.; validation: V.M., S.R., S.D.B.; M.F., formal analysis: S.D.B., and V.M.; investigation: S.D.B., and S.R.; writing-original draft preparation: S.D.B.; writing-review and editing: G.P., L.M., V.M., and S.D.B. All authors read and approved the final version of manuscript.

## Ethics approval and consent to participate

All participants completed the informed consent form to the study protocol which was approved by the Ethics Committee of the German Society of Psychology, UL 072014.

## Supplementary Material

### 1 Cognitive Tests

#### 1.1 Episodic Memory (EM)

##### Object Location task (Object)

In this task, sequences of 12 colored photographs of real-world objects were displayed at different locations in a 6-by-6 grid. After presentation, objects appeared at the side of the screen and had to be moved to the correct locations by clicking on the objects and the locations with the computer mouse. One practice trial and two test trials were included. The sum of correct placements across the two test trials is used as the manifest variable.

##### Face–Profession task (Face)

This task assesses associative binding on the basis of recognition of incidentally encoded face–profession pairs. During the study phase 45 face–profession pairs were each presented for 3.5 s on the computer screen and the participants had to indicate via button presses whether the faces matched the profession or not. After a 3-min delay between study and test phase 54 face–profession pairs consisting of 27 old pairs, 9 new pairs, and 18 newly arranged pairs were presented (in newly arranged pairs the shown face is the same, but is associated with a new profession). The participants were asked to decide whether they had seen a given face–profession combination before or not and to rate the confidence of their decision on a three-point scale ranging from 1 = not sure to 3 = very sure. Recognition memory for the rearranged face–profession pairs (hit minus false alarms) served as the manifest variable in the model.

#### 1.2 Working Memory (WM)

##### Number-N-Back (Number)

Three one-digit numbers (ranging from 0 to 9) were presented sequentially in three cells situated horizontally followed by the next sequence of three digits. This cycle was repeated 30 times. In each cycle, two-choice decisions on whether the current stimulus matched the stimulus shown three steps earlier had to be made. Four practice trials including 30 runs were followed by 6 test trials with 30 runs. Subjects made their decision via button-box presses with their left and right index fingers (Schmiedek et al. 2010).

##### Letter Updating (LetterU)

In this task subjects were presented with 7, 9, 11, or 13 letters in a sequence. Once a sequence stopped, subjects had to report the last three letters in correct order by pressing buttons on the button box corresponding to A, B, C, and D.

##### Spatial Updating (Spatial)

In each block of this task, a display of two or three 3-by-3 grids was shown for 4 s. In each of these, one blue dot was presented in one of the nine locations. Those two or three locations had to be memorized and updated according to shifting operations that were indicated by arrows appearing below the corresponding field. The presentation time of the arrows was 2.5 s with an inter-stimulus interval of 0.5 s. After six updating operations, the two or three grids reappeared and the resulting end positions had to be clicked on. After ten practice blocks with memory loads of two and three grids, ten test blocks with load two and three were conducted and used for scoring. The average percentage of correct placements was used as one of the manifest variables for the working memory (WM) factor (Schmiedek et al. 2010).

#### 1.3 Fluid Intelligence (Gf)

##### Figural Analogies (Analog)

Items in this test followed the format "A is to B as C is to ?". One figure pair was presented in the upper left part of the screen and a single figure was shown beside it. Participants had to use the same rule as the one applying to the complete figure pair to choose one of the five alternative responses presented below. Subjects entered their response by clicking on one of the five alternatives with the mouse. Before the test phase, instructions and three practice items were given. The test phase was terminated when subjects made three consecutive errors, when they reached the maximum time limit (10 min), or after they had worked on the last test item. Items were ordered by difficulty (Lindenberger et al. 1993).

##### Practical Problems (Problem)

This task consisted of 12 items depicting everyday problems such as the times in a bus schedule, instructions for medication, a warranty for a technical appliance, a rail map, as well as other forms and tables. For each item, the problems were presented in the upper part of the screen, and five alternative responses were shown in the lower part. Subjects responded by clicking on one of the five alternatives with the computer mouse. A single practice item was provided. The test phase was terminated when subjects made three consecutive errors, or when they reached the maximum time limit of 10 minutes, or after they had answered the last test item. Items were ordered by difficulty (Lindenberger et al. 1993).

##### Letter Series (Letter)

The task consisted of 22 items. Each item contained five letters followed by a question mark (e.g., c e g i k ?). Items were displayed in the upper half of the screen, and five response alternatives were presented in the lower half. Items followed simple rules such as +1, –1,+2, or +2 +1. Subjects entered their response by touching one of the five answer alternatives. The score was based on the total number of correct responses. Instructions and three practice items were given before the test phase. The test phase was terminated when subjects made three consecutive false responses, when they reached the maximum time limit (6 min), or after they had answered the last item of the test. Items were ordered by difficulty. Sample items were used with respect to tests related to speed, reasoning, and knowledge (Lindenberger et al. 1993).

#### 1.4 Processing Speed / Comparison Task (Speed)

For *the numerical version (number)* of the comparison task, two strings of five numbers each appear on the left and right of the screen, with participants having to decide as quickly as possible whether both strings are exactly the same or different. If different, the strings differ by just one number. Number strings are randomly assembled using digits 1–9.

*The verbal version (verbal)* of this task is equivalent to the numerical one, using strings of five consonants.

##### Figural version (figure)

Two “fribbles” - the three-dimensional colored objects consisting of several connected parts, are shown to the left and right of the screen, with participants having to decide as quickly as possible whether the two objects are exactly the same or different. If different, the objects differ with respect to one part. The Fribble images in this task are courtesy of Michael J. Tarr, Brown University, http://www.tarrlab.org/). In the session, two trials of 40 items were included for each of the verbal, numerical, and figural tasks.

**Supplementary Figure 1.**
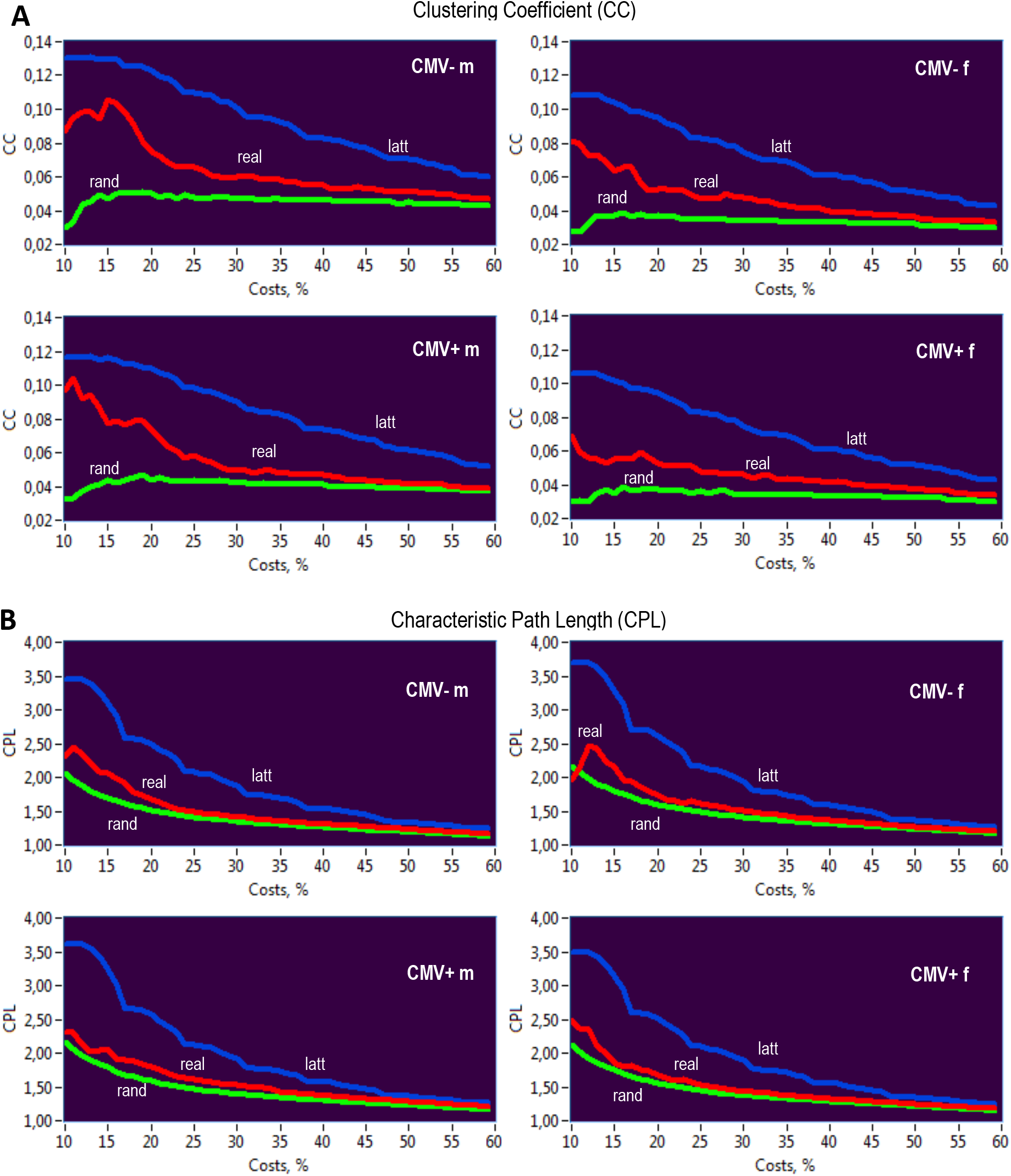
(**A**) *CC* is greatest in lattice networks (blue) and lowest in random networks (green), whereas *CC* for the real networks (red) is in-between. In contrast, (**B**) *CPL* is shortest in random and longest in lattice networks, while the real networks are in-between. CMV, Cytomegalovirus; CMV^−^ m, CMV-seronegative men; CMV^+^ m, CMV-seropositive men; CMV^−^ f, CMV-seronegative women; CMV^+^ f, CMV-seropositive women.

**Supplementary Figure 2.**
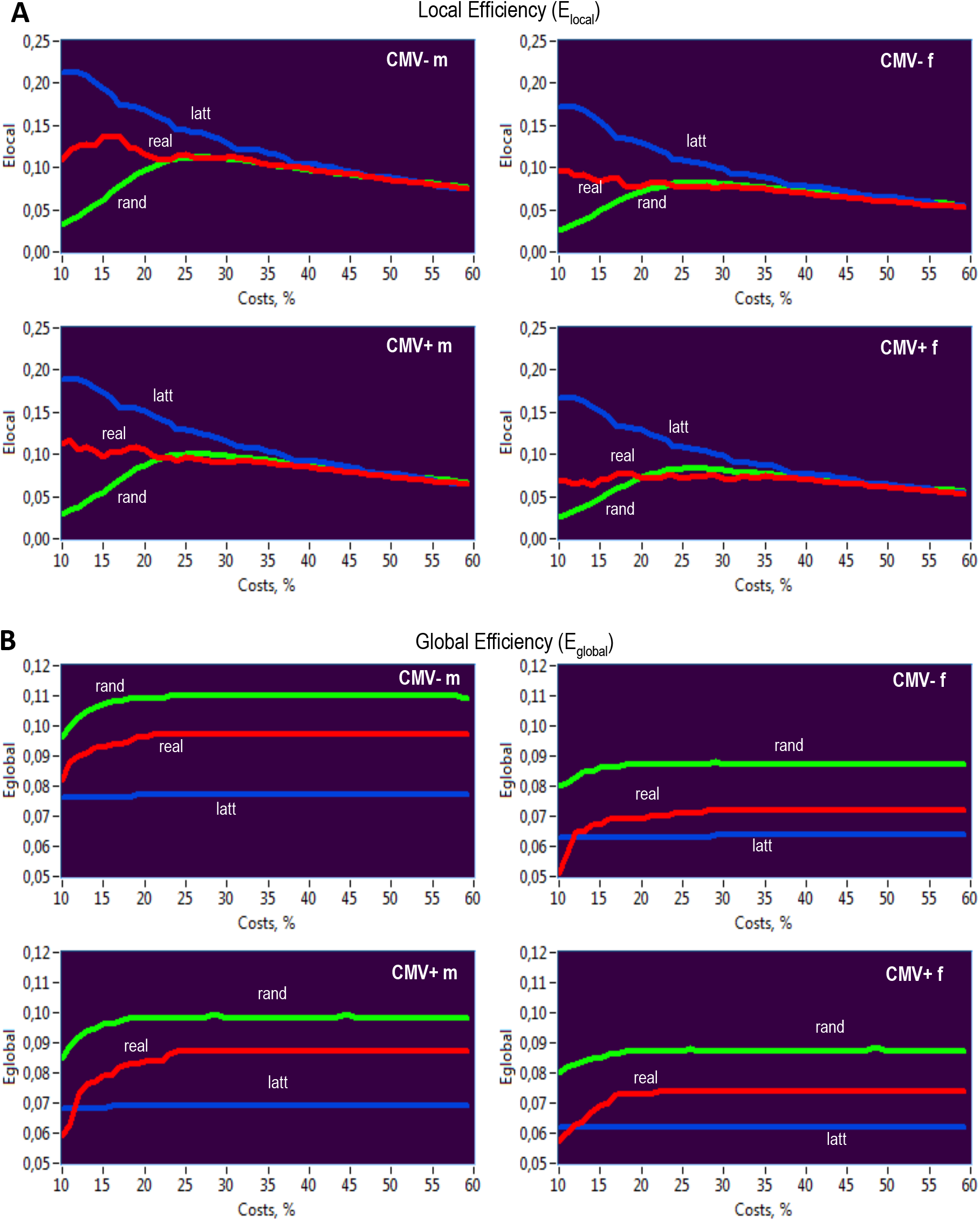
(**A**) Local efficiency was highest in regular networks (at least for the cost levels under 45%) and lowest in random networks (at least for the cost levels under 20%), while (**B**) global efficiency was highest in random (green) and lowest in lattice (blue) networks practically for all levels of wiring costs, with real (red) networks were always in-between. CMV, Cytomegalovirus; CMV^−^ m, CMV-seronegative men; CMV^+^ m, CMV-seropositive men; CMV^−^ f, CMV-seronegative women; CMV^+^ f, CMV-seropositive women.

